# Reconstruction of Average Subtracted Tubular Regions (RASTR) Enables Structure Determination of Tubular Filaments by Cryo-EM

**DOI:** 10.1101/774802

**Authors:** Peter S. Randolph, Scott M. Stagg

**Affiliations:** Institute of Molecular Biophysics, Florida State University, Tallahassee, FL 32306, USA; Department of Chemistry and Biochemistry, Florida State University, Tallahassee, FL 32306, USA

## Abstract

As the field of electron microscopy moves forward, the increasing complexity of samples being produced demand more involved processing methods. In this study, we have developed a new processing method for generating 3D reconstructions of tubular structures. Tubular biomolecules are common throughout many cellular processes and are appealing targets for a variety of biophysical research. Processing of tubules with helical symmetry is relatively straightforward for electron microscopy if the helical parameters are known, but tubular structures that deviate from helical symmetry (asymmetrical components, local but no global order, etc) present myriad issues. Here we present a new processing technique called Reconstruction of Average Subtracted Tubular Regions (RASTR), which was developed to reconstruct tubular structures without applying symmetry. We explain the RASTR approach and quantify its performance using three examples: a simulated symmetrical tubular filament, a symmetrical tubular filament from cryo-EM data, and a membrane tubule coated with locally ordered but not globally ordered proteins.

## Introduction

Recent technological innovations have significantly increased the efficiency and resolution of structures determined by cryogenic electron microscopy (cryo-EM), causing a proliferation in the number of structures resolved with ever-increasing resolution^1–4^. As many of the “low hanging fruit” structures have been determined, studies have shifted toward increasingly complex challenges, pushing the capabilities of current processing software, and many requiring novel methodologies. Here we present a new approach called Reconstruction of Average Subtracted Tubular Regions (RASTR) which is designed to examine asymmetric architectural features of tubular structures.

Tubular structures are common throughout all domains of life. Tubule architecture is advantageous for a variety of functions. The large surface area to volume ratio can provide a platform for scission/fusion, gated compartments, reaction surfaces, etc^5–7^. In addition, tubules can form transport networks, allowing directed transport to distant structures/areas^8,9^, and be used for motility^10,11^. Tubules are usually formed by one of three processes: 1) self-assembly of lipid tubules when the appropriate lipids or small molecule(s) are present^12^, 2) deformation of membranes through protein insertion^12,13^, or 3) helical arrangements of proteins (polymers)^14^.

Helical polymers (microtubules, actin/myosin, pila, etc) have been studied extensively and are central to myriad processes across all domains of life, such as trafficking routes^8,9^, motility^10,11^, mechanical support^15,16^, muscle force^17,18^, etc. Helical polymers have been targets for electron microscopy since the technique’s inception^19^. Resolving the structures of simple helical complexes can be relatively straightforward^20^, but tubular structures lacking in helical symmetry, with bends/imperfections, or asymmetric components still present numerous issues.

The first resolved structure containing helical architecture is the Bacteriophage T4 tail^19^. Since then improvements in technology have allowed for a drastic increase in resolution, with a recent helical structure being determined to 1.9 Å^21^ resolution. Originally, the structures of helical tubules have been solved by treating the tubules as helical crystals using the Fourier-Bessel approach^22^. However, Bessel overlap can make unambiguous determination of the helical symmetry for these specimens challenging, especially for tubules of large diameter^24^. As the field has advanced, multiple techniques have been developed to deal with structures whose architecture does not conform to perfect symmetry. One popular approach is Iterative real-space helical reconstruction (IRSHR)^23^. IRSHR separates the helical filament into small segments which are then treated as single particles. The segments are aligned, and a 3D model is generated before helical symmetry is determined and then applied. This allowed IHRSR to deal with a variety of issues that plagued the Fourier-Bessel approach for less than ideal samples. Utilizing helical symmetry has many advantages; each micrograph contains all possible views of the particle (unlike 2D crystals), easily aligned (compared to single particles), and the particles will be averaged to get to a higher resolution. Unfortunately, this process can be a double-edged sword, as utilizing helical symmetry increases the resolution for the helical portions, but blurs features that do not follow the helical symmetry. Asymmetric features such as myosin heads, decorations, etc. are averaged out of the final structure.

There are many examples of proteins that interact with helical polymers, actin-binding proteins by themselves run the range from A (a-actinin^25^) to Z (Zyxin^26^). When proteins decorate tubules, they can form complexes that are asymmetrical, preventing the use of helical symmetry for resolving the architecture. These decorations contain many pathologically interesting targets as mutations causing improper formation of these complexes can lead to a variety of disease states including but not limited to chronic myelogenous leukemia (CML) (Abl)^27^, elliptocytosis (Protein 4.1)^28^, and inhibition of tumor suppression (p53)^29^.

Tubules can also form by perturbation of lipids bilayers through either small molecules/unique lipid headgroups^30,31^ or insertion of proteins^13,32,33^. Lipids commonly form bilayers, micelles, or vesicles, but the insertion of proteins chains (usually α helices) causes an increase in the surface area of one leaflet of the bilayer, forcing the architecture to deform into a variety of shapes including tubules^34^. Lipid tubules make up several intracellular organelles, such as the endoplasmic reticulum and trans-Golgi network. Their high curvature and large surface area to volume make them ideal for protein accumulation and inter-organelle transport^7^. Many proteins that perturb membrane structure form an ordered array (similar to a 2D lattice), which allows for uniform dimensions of the tubule and as a benefit allow easier reconstruction of the structure. Some proteins (Sar1 tubules^33^) appear to only form small areas of local order, which limits the methods of resolving the architecture.

We have developed a technique for upweighting a section of the surface of tubular specimens in order to resolve their structures without applying symmetry. This enables structure determination of decorations on tubules or filaments and has the potential to enable structure determination of tubules with ambiguous symmetry. Our technique, called reconstruction of average subtracted tubular regions (RASTR), combines multiple electron microscopy processing tools to isolate a section of the tubule and upweight one side. Once sections are isolated and upweighted using RASTR, they are then treated as single particles and aligned and reconstructed. Here we present the RASTR process, and reconstructions of a helical filament from ideal and experimental data, without imposing symmetry.

## Materials and Methods

### GalCer Tubes

Lipid nanotubes with a uniform diameter (25 nM) were constructed with molar equivalents of D-galactosyl-β-1,1’N-nervonoyl-D-erythro-sphingosine (GalCer) and a lipid mix (55% DOPC, 35% DOPS, and 10% cholesterol by molarity). GalCer and lipids were dissolved in chloroform which was dried under argon and rehydrated in a minimal salt buffer (25 mM HEPES pH 7.2, 1 mM Mg(OAc)_2_) to a final concentration of 0.8 mg/ml by vortexing. Tubes were assessed through negative stain TEM (2% Uranyl acetate) on a CM120 Biotwin and then cryogenically frozen with a FEI Vitrobot on c-flat holey carbon grids (2/2 300).

### Generating ideal sample data

Sample data was generated from the VipA/VipB model from Kudryashev, et al (EMPIAR-10019)^8^. The VipA/VipB filament was chosen because of the high-resolution model, the readily available experimental data, and the high degree of symmetry. The pdb model (3J9G) was elongated to span the length of the box in pymol^35^ utilizing symmetry and alignment, then a 3.5 Å map was created with pdb2mrc from the EMAN suite. The map was projected using a sample star file containing ctf’s from the VipA/VipB experimental data, randomized phi, theta = ± 90° (generate tubes with opposite polarities), psi = 90°, and x,y shifts = 0, creating a stack with box size 240×240 and 2 Å per pixel. No noise was given for the generated ideal data. Both the experimental data (available from EMPIAR) and the generated ideal sample data were used.

### Sar1…GalCer Tubes

Sar1 was expressed and purified in the same manner as Hariri et al.^33^. Sar1…GalCer tubes were produced by incubating GalCer tubes with 1.5 mM GTP (or 5’-Guanylyl imidodiphosphate (GNPPNP)), and 0.7 µM Sar1 for 2 hours at 42 C. Tubes were assessed through negative stain TEM (2% Uranyl acetate) on a CM120 BioTwin and then cryogenically frozen with a FEI Vitrobot on c-flat holey carbon grids (2/2, 300).

### Data collection and processing

All data collected in-house was done on a Titan Krios 300kV with a DE64 detector in integrating mode. Frames were collected in the Appion/Leginon^36,37^ environment, aligned with MotionCor2^38^, and CTF estimated with both GCTF v1.06^39^ and CTFFIND v4^40^. The final CTF values were chosen based on what had the highest resolution agreement between the estimated and measured CTF. Tubes were manually selected along the long axis of the filaments, though no symmetry was supplied. Particles were roughly aligned during stack creation so that the filament axis aligned with the Y axis.

### Post-processing and validation

A variety of aligning and classifying programs were used. cisTEM^41^ was used for most of the 3D alignment and refinement, and cryoSPARC^42^ for the majority of 2D alignment and classification. RELION^43^ was used as an alternative for 2D classification and 3D refinement. The RASTR program uses RELION subtraction, so all models are run through RELION 3D refinement without alignment prior to RASTR processing. The final map was masked to just the upweighted region and sharpened using Phenix Autosharpen Map^44^. Map validation was done with Phenix Mtriage^45^ and ResMap^46^.

### RASTR Methodology

The objective of RASTR is to isolate discrete areas on the surface of the tubule. To properly isolate these surfaces, we generated a methodology which consisted of distinct steps that we wrapped in a program called RASTR. Here we present this methodology.

A. During the first step of the RASTR process, tubules were aligned, and their Euler angles randomized about the azimuthal axis which generated an azimuthal-averaged (AA) model (Fig. 2A). We typically used cisTEM and an in-house python script to randomize phi for this step due to the ability to control the Euler search parameters with cisTEM, but theoretically any 3D refinement software should suffice. After randomization, the tubule was reconstructed generating an AA model. Because subtraction was carried out in RELION, the model was run through relion_refine without alignment to set the weighting for the RELION suite. The input files necessary for RASTR are the original micrographs, AA model, and the phi-randomized star file.
B. First RASTR masked out a region of interest in the AA models (generating *n* models with the masked region at an angle of (360/*n*)° (Fig. 2B). A numpy array was created with the same dimensions as the input model and populated with values of 1 except for a sphere centered at the given *x* value and of radius *r*, which contains values of 0. This array was outputted as a 3D model (the mask) and low-pass filtered to generate a gaussian edge (size of gaussian padding is decided by the user). The mask was multiplied by the input AA model, resulting in a tubule with a sphere masked out (model-phi000.mrc). This was repeated with a new sphere rotated in increments of (360/*n*)°, generating (Model-phi(360/*n*).mrc, Model-phi(2* (360/*n*).mrc, …, Model-phi((*n*-1)* (360/*n*).mrc). By generating multiple models rotated about the azimuthal axis, we were able to capture the entire circumference of the tubule, as well as generating overlap for alignment. Subtraction near the edges of the box was found to create artifacts. The combination of providing proper coverage (inclucing overlap for symmetrical particles) and avoiding the borders means that overlap on the azimuthal axis should be generated during initial stack creation.
C. RASTR generated and subtracted projections of the masked AA models from the original micrographs (upweighting the regions of interest relative to the background) (Fig. 2C), also subtracting signal from the opposite side of the tubule which was projected in the same area. Relion_project was used to project and subtract the masked tubules (model-phiXXX.mrc) from the original micrographs. The initial star file for the AA model provides the Euler angles, shifts, and ctf. Projection and subtraction was done for each model, generating an upweighted stack for each.
D. The upweighted regions were then further masked (Fig. 2D), leaving only the area of interest, which had a small amount of signal removed during subtraction which should correspond to the opposite face of the tubule, allowing isolation of one face. Using the provided Euler angles from the supplied star file, the center of the masked area was tracked from the masked model to the upweighted stack using the rotation matrix below, where Z_1_ is phi, Y_2_ is theta, and Z_3_ is psi. The rotation matrix generated a new (*x,y,z*) but the *z* was dropped because the model is being projected on a 2D plane. (*x,y*) were offset by the provided *x* and *y* shifts. A new stack was created masking the pixels outside the upweighted area with the mean value of the original particle image.

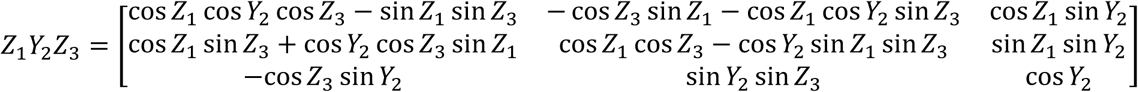 The AA model was a smoothed estimate of the face, so while the signal was reduced, the information will remain and generates some noise to the final model but does not appear to affect resolution as it generates a degraded structure outside the upweighted region which can be masked out. If desired, the area of upweighting can be extracted into a smaller box, though this process is optional, and we have found it increases the noise in the final model. For RASTR arguments see Supplemental Section 1.
E. In the last step, the star files of the masked particles were concatenated into a single star file (directing to the various stacks). Relion_preprocess was run to create a single stack and star file and relion_reconstruct was used to generate an initial model for alignment and to spot-check proper upweighting/masking and alignment. After RASTR processing the RASTR particles were treated as a stack of single particles, with some caveats. As the particles are removed from the surface of a tube they are limited in their degrees of freedom for alignment. If the psi and theta alignment were perfect before RASTR processing, then only phi and y-shift would need to be refined. This was obviously not the case, but psi, theta, and x-shifts should be limited to small changes, and any large deviations can be dismissed as misalignment.

**Figure 1.**
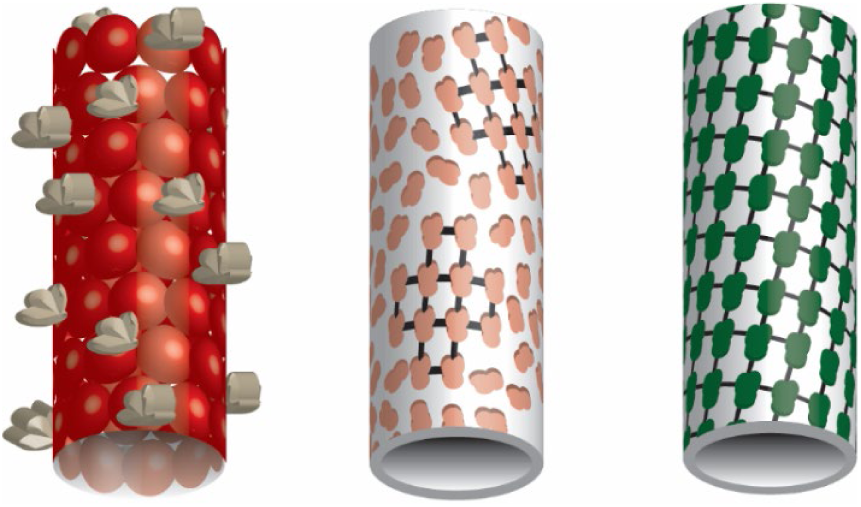
Tubule examples. From left to right, a tubule f i lament made f rom ordered repeats of protein with decorations (left). Membrane tubule with decorating particles with areas of local order (no global order) (middle). Membrane tubule decorated with particles with global order (2D crystal) (right).

**Figure 2.**
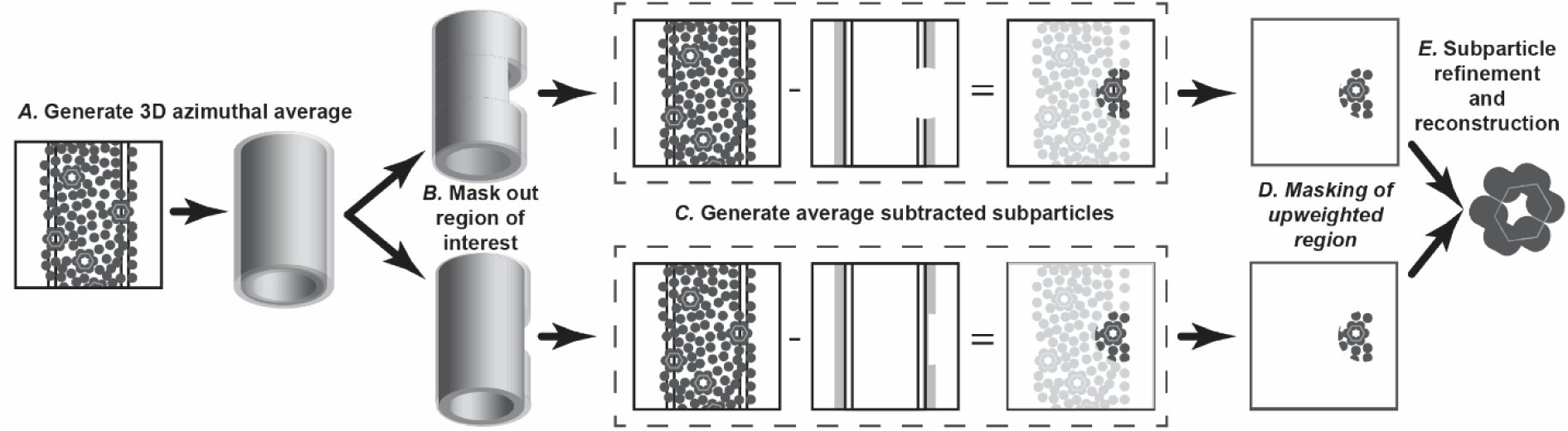
RASTR procedure schematic. **A** An azimuthal-averaged (AA) model was generated f rom the tubule data. **B** The region of interest was masked out in the aa model, either the tubule wall and decorations (top) or just the decorations (bottom). **C** The masked aa model was projected and subtracted f rom the original particles, upweighting the region of interest. **D** The region of interest was masked. **E** The individual regions are aligned and refined as single particles.

## Results and Discussion

### Bare GalCer tubules

The success of RASTR is highly dependent on the effectiveness of the model subtraction. To test subtraction, we collected undecorated GalCer tubes (See Supp. Table 1 for data collection statistics), aligned in cisTEM (Fig. 3A-B) and an AA model was generated (Fig. 3C). The AA model was subtracted from the original particles (Fig. 3D-F), in this case without specifying a region for upweighting so that the complete tubule could be subtracted.

**Figure 3.**
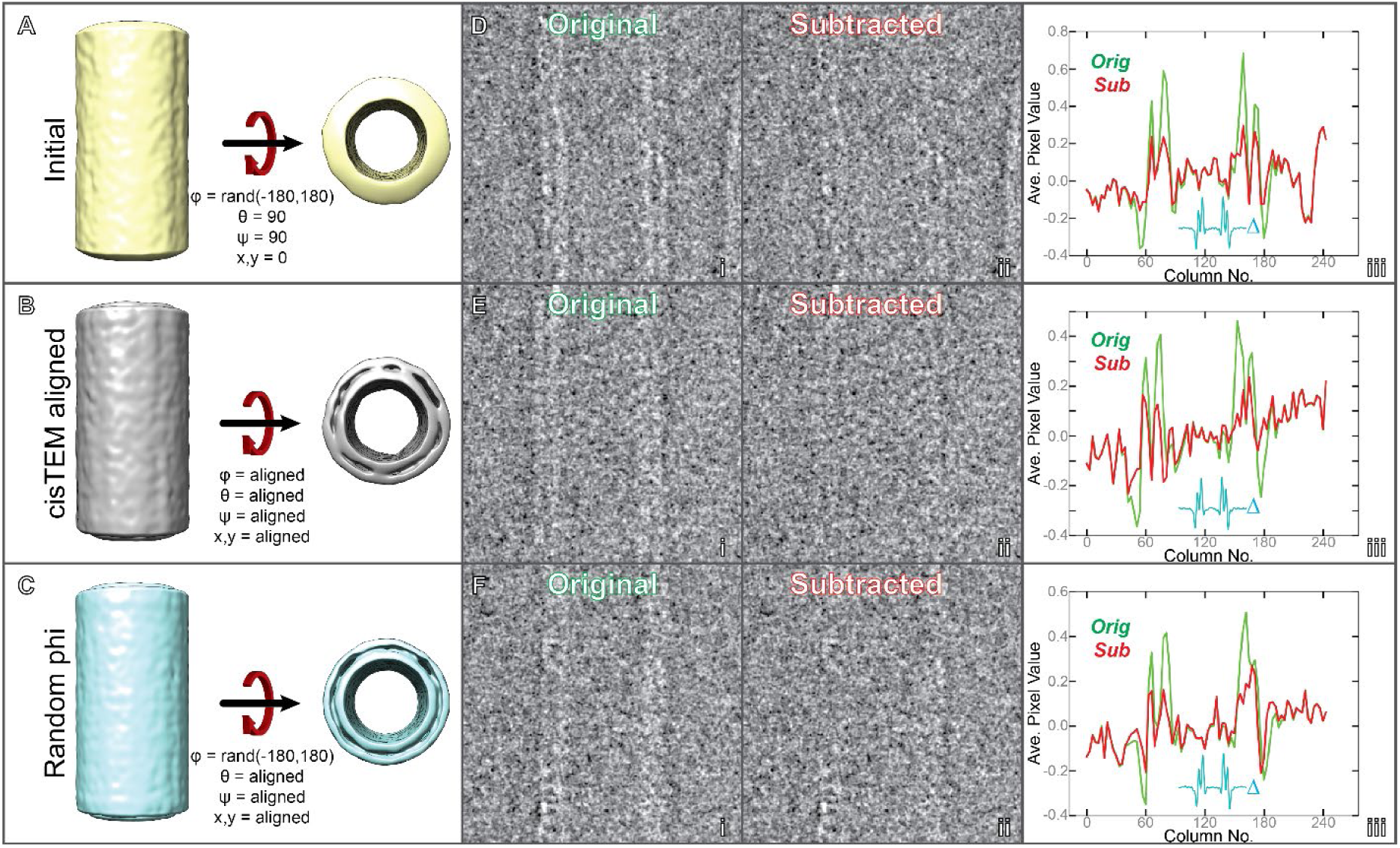
Subtraction of bare Gal Cer tubules. **A** Gal Cer tubes were roughly aligned vertically in two dimensions during stack creation then φ was randomized between - 180° and 180°; and θ, Ψ were set to 90° for reconstruction. **B** All Eulers and shifts were aligned in cis TEM. **C** φ was once more randomized between −180° and 180°, creating an azimuthal-averaged model. **D**,**E**,**F** Examples of undecorated Gal Cer tubule particles before (i) and after (i i) subtraction with relion_project.(iii) The average pixel value of each column (excluding a ten pixel border along the edges of the image). Original image (green) and subtracted (red), with the difference inlayed in cyan (Δ).

The azimuthal-averaged model was subtracted from the original particles using relion_subtract. Three representative samples are shown in Fig. 3D-F. Each particle has features that illustrate the pros and cons of our current method of subtraction. Particle D was generally straight, has a distinct bilayer, but had a slight deviation in diameter and a kink at the top. Particle E was an ideal particle, as it was straight and has a distinct bilayer. Particle F was straight, but the right bilayer was not distinct. The average pixel value of each row (with a ten pixel border to remove artifacts from normalization) is presented in Fig. 3D,E,F(iii) to provide an objective measure of the subtraction. The subtraction worked well for all cases but was most successful on particle E, with the average values almost reduced to noise, while remnants can be seen in both particles D and F. The most consistent subtraction was the center sections of the particles, away from the edges. It should be noted that the diameters of the tubes have the potential to have a large effect on the subtraction. If the particles’ diameters deviate from the AA model, then instead of subtracting features the subtraction will create artifactual negative densities. For particles with a consistent diameter (including most tubule filaments), one model suffices, but for tubules that vary in diameter the particles would need to be classified before alignment and distinct azimuthal-averaged models generated for each class.

### Ideal VipA/VipB

After the individual sections of RASTR were tested, we tested RASTR on simulated data in order to assess the ability of the approach to faithfully reconstruct a biological filament. Ideal simulated images of tubular filaments were created using the VipA/VipB pdb model^8^ (see Methods), with the original particle stack containing 3550 particles, equivalent to 426,000 ASUs (Fig. 4D, left). Reconstruction of the projected particles regenerated the original ideal map (Fig. 4A). An azimuthal-averaged model was created (Fig, 4B) and processed in RASTR, upweighting a radius 50 spherical section, 50 pixels from the center of the box on the *x*-axis and repeated at increments of 15° (360°/24) around the azimuth (Fig. 4C). The final stack contained 85,200 sub-particles (Fig. 4D, right), equivalent to 150,000 unique ASUs. Particles were reconstructed using the azimuthal-averaged Eulers and shifts (offset by 15° increments) (Fig. 4E), which recaptured the missing area from the original subtraction model (Fig. 4F). Using cisTEM, we were able to successfully align the ideal RASTR VipA/VipB data, resolving the original structure only in the region we had upweighted (Fig 5). Since there was residual density in the images due to the imperfect subtraction of the AA model, there was elongated and diffuse density in the 3D reconstruction in the regions outside of the upweighted region of the tubule. (Fig. 5B). Local resolution calculated in ResMap confirms this result, with the upweighted region showing a resolution around 4Å, which rapidly deteriorates as one goes farther from the region until all density is lost (Supp. Fig 2,3). The RASTR aligned map recaptures the details of the original model, from the core filaments to the extended α-helices and loops (Fig. 5D). The reconstruction of the ideal data from VipA/VipB provided a blueprint for how to treat filaments in RASTR. The lack of definition in the areas outside the upweighted section, meant a mask was necessary or the reconstruction began to degrade. Another issue we found was during initial alignment the particles tended to bunch around the starting phi angles producing an extremely low-resolution map. Allowing the other angles freedom to align only exacerbated the issue as the particles were still bunched but they would diverge from reasonably allowed translational and rotational alignments for the tube resulting in a junk reconstruction. Fixing the alignment to just phi and y-shifts until the particles spread out around the azimuthal axis and had an initial alignment then freeing the other angles to refine resulted in a significantly better reconstruction (See Euler angle distribution Supp. Fig 1).

**Figure 4.**
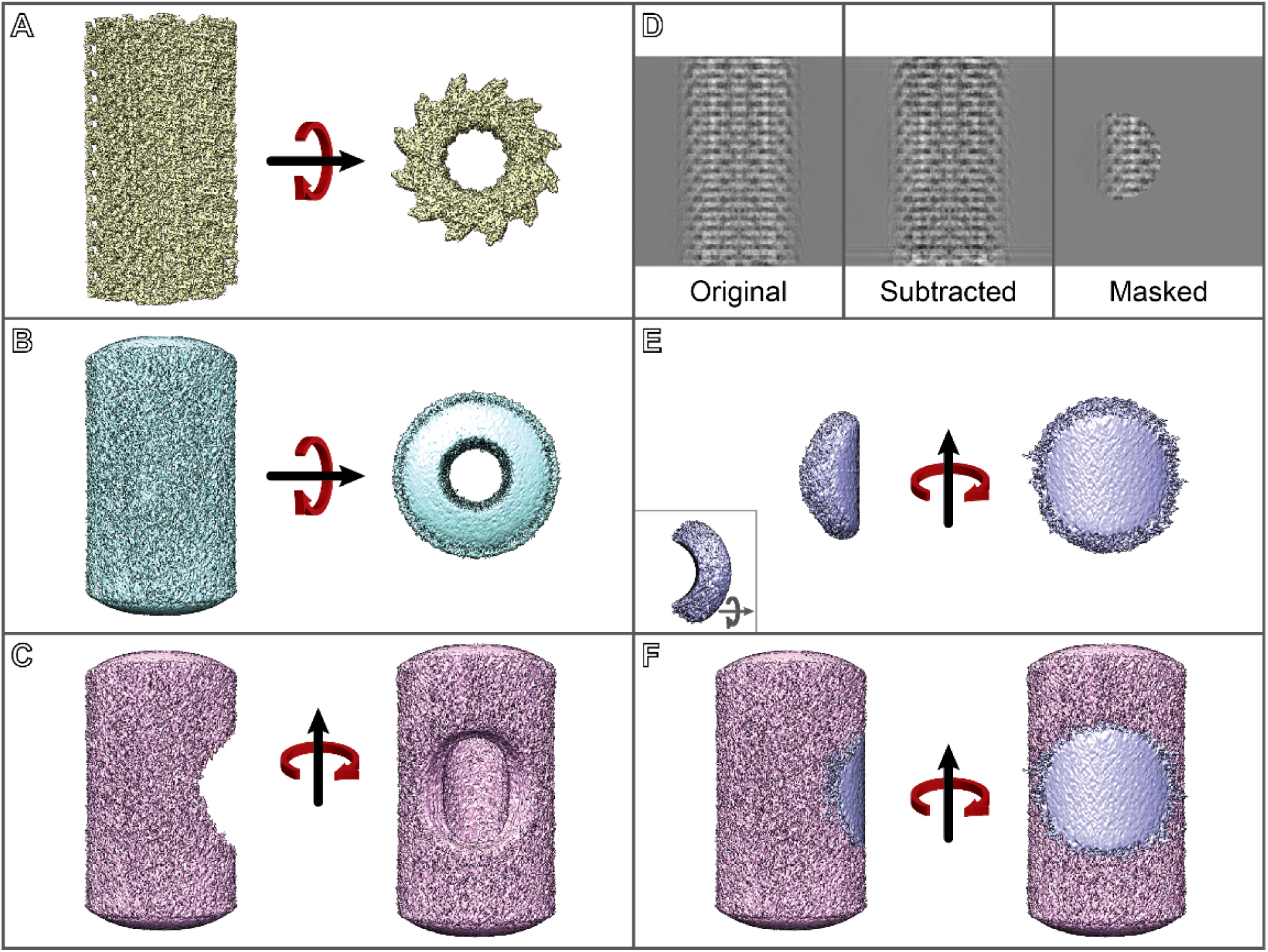
RASTR summary using ideal Vip A/ Vip B data. **A** Extended VipA/ VipB model generated f rom pdb 3J 9G. **B** Azimuthal-averaged VipA/ VipB model. **C** Model to be subtracted f rom the particles with the section to upweighted removed. **D** Particle progression through RASTR, f rom left-to-right: original particle, subtracted, and then masked. **E** Reconstruction of the final RASTR particles using the Eulers and t ranslational shifts f rom the azimuthal-averaged model. **F** Overlay of the model to be subtracted (**C**) and reconstruction of the RASTR particles (**E**).

**Figure 5.**
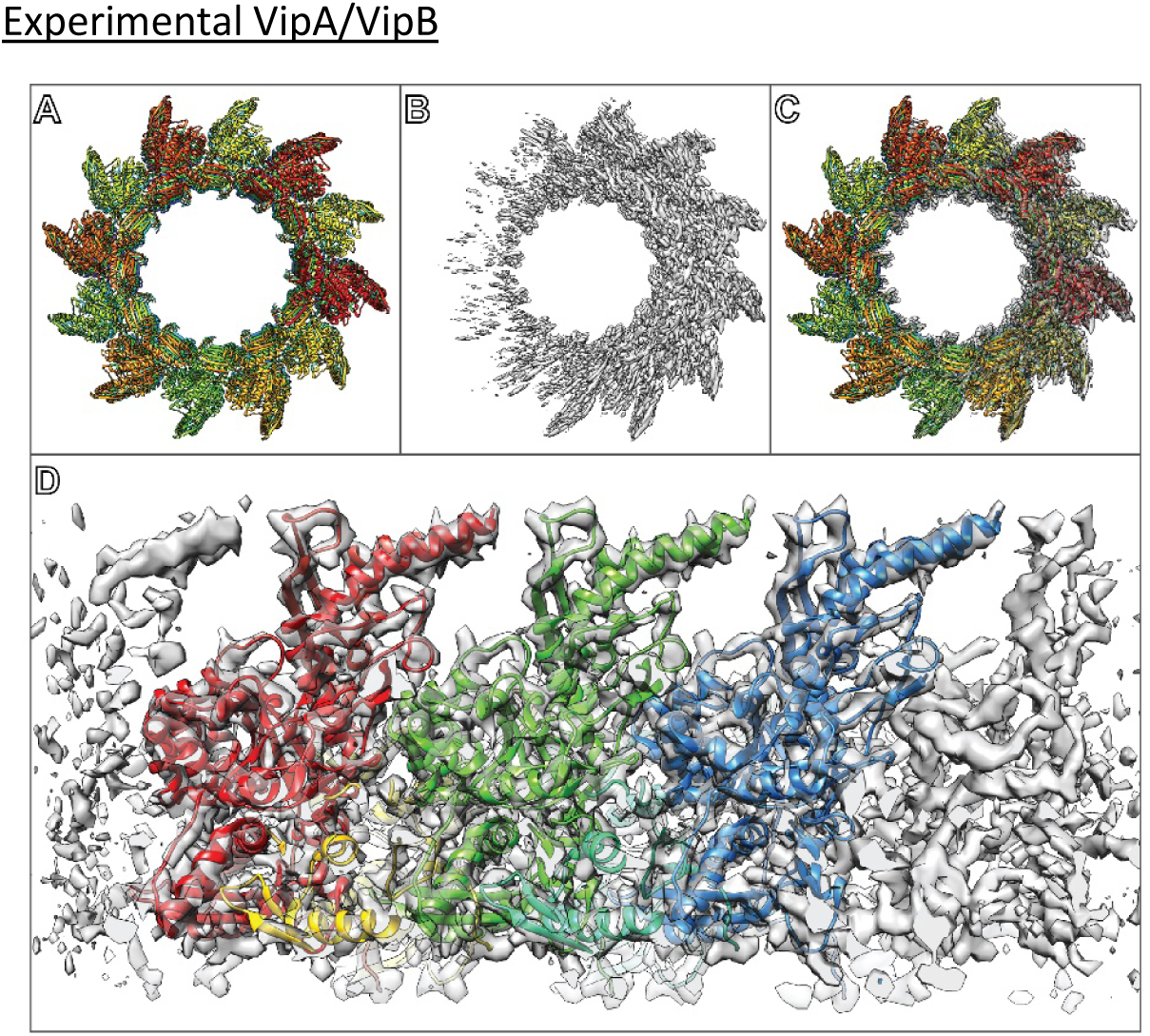
Ideal data (generated Vip A/Vip B) filament alignment. **A** Original VipA/ VipB pdb model (3J 9G), extended along the azimuth. **B** Aligned map of RASTR VipA/ VipB particles. The middle right section of the aligned map resolved to a high resolution, which degrades as the angle away f rom the middle right increases. **C** Overlay of (**A**) and (**B**). **D** Middle right section of the aligned map f i t the original data, showing secondary structure.

### Experimental VipA/VipB

Given the success with RASTR on simulated data, we applied the technique to real experimental cryo-EM data. Experimental VipA/B cryo-EM data^8^ are available on the EMPAIR database, and these were obtained from EMPIAR and processed in RASTR. The VipA/VipB structure was originally resolved using Iterative Helical Real Space Reconstruction (IHRSR) to a resolution of 3.5 Å^8^. For RASTR, the filaments were manually picked, and 480 pixel stacks created with 1 Å/pixel. The helical step during stack creation was only 50 Å, generating significant overlap between particles, which was necessary for RASTR processing. The VipA/VipB filaments were processed in RASTR, upweighting a spherical section that was 100 pixels from the azimuthal average (centered on the highest density of VipA/VipB) and with a 100 pixel radius. cisTEM was used to align the RASTR processed VipA/VipB data resulting in a final map with a reported resolution of 4 Å which is lower than the reported IHRSR structure, however the reconstruction of the RASTR processed particles produced a map with improved features compared to the original helically aligned one.

As with the ideal data, RASTR processed data produced a high-resolution map in the upweighted sphere but with deteriorating quality outside of the upweighted sphere (Fig. 6B). Since the only valid part of the 3D map was inside the upweighted region, all subsequent analysis were limited to that section (Fig. 6B, highlighted). The threshold was normalized between the original IHRSR map and the RASTR reconstructed map by zoning the maps around a single VipA/VipB unit (one VipA chain and one VipB chain) and setting the threshold so the enclosed volumes were equal. Both maps produced a high enough level of detail, including side chain density, for the architecture from the inner surface (Fig. 6C) through the buried residues (referred to as Domain 1 and Domain 2). Two exposed α-helices (VipA H4 and VipB H1) on the outer surface of the filament (Domain 3), showed more isotropic density using the RASTR process (Fig. 6D). In addition, we were able to recapture a section of unmodeled density that was observed in the original helical reconstruction (Fig. 7, maps filtered to 7.5 Å). This density had higher occupancy for the RASTR map and resolved as two distinct elongated blobs running parallel to each other, possibly two α-helices.

**Figure 6.**
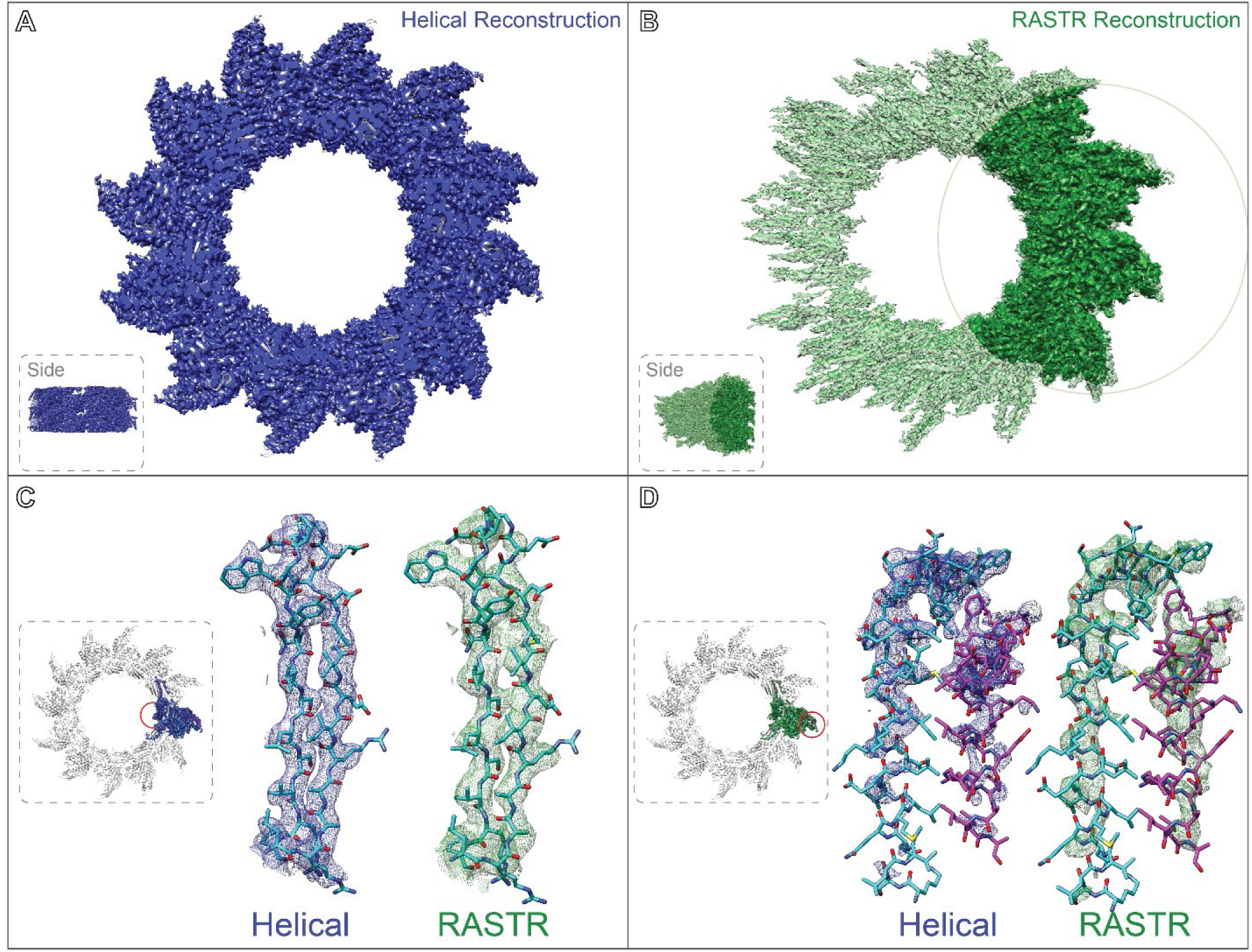
Experimental VipA/ Vip B sheath reconstruction. **A** Original reconstructed map aligned using IHRSR (blue). **B** RASTR processed reconstruction with upweighted area highlighted (green). **C** Domain 1 β-sheets f rom the inner surface. Both maps had good detail, with the original model generated f rom the IHRSR alignment fit in the new RASTR map. **D** Two outer surface α-helices f rom Domain 3 had less coverage in the IHRSR map (blue) than the RASTR map (green), though the improvement was slight.

**Figure 7.**
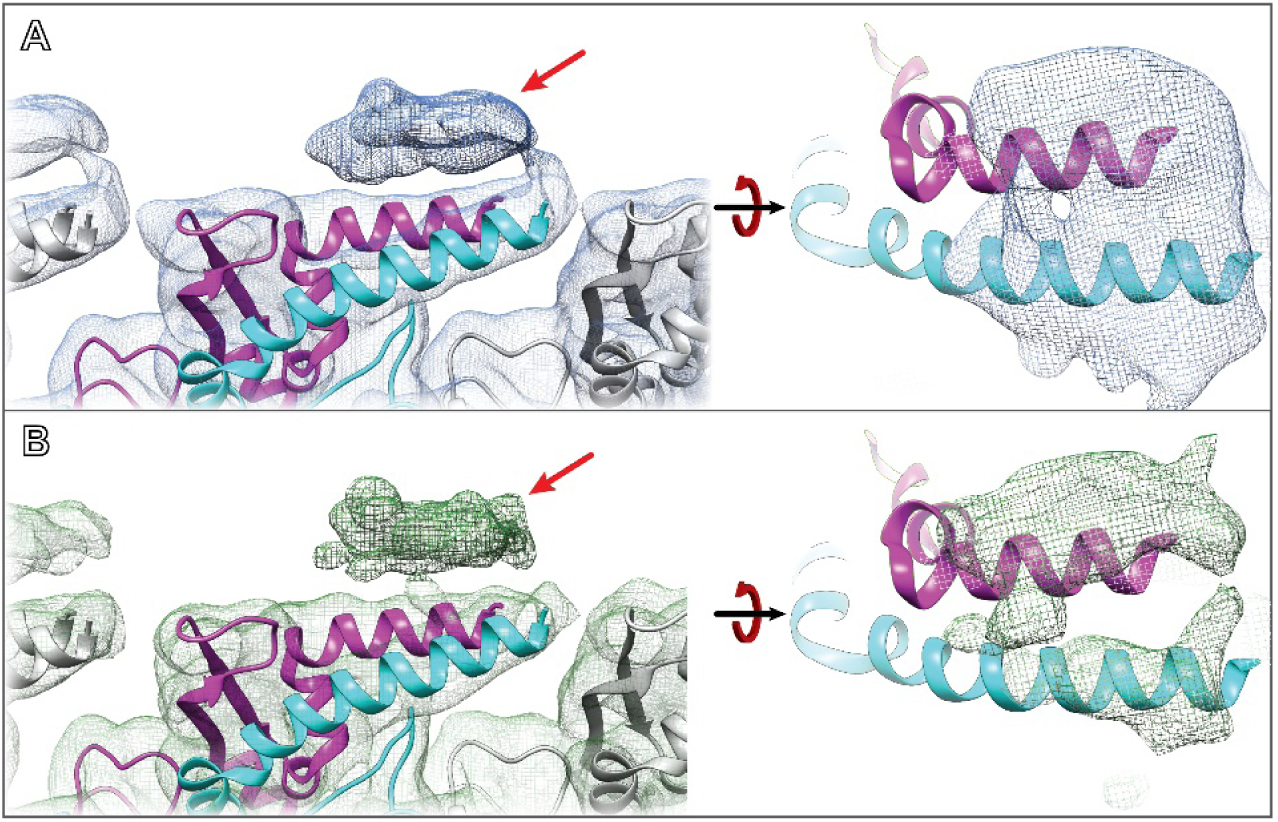
VipA/ VipB sheath unmodeled density. **A** Original map f i l tered to 7. 5 Å highlighting the unmodeled density identified after alignment (left) with top view (right). **B** RASTR map f i l tered to 7. 5 Å exhibiting the same density (left) with top view showing two distinct oblong densities running parallel.

**Figure 8.**
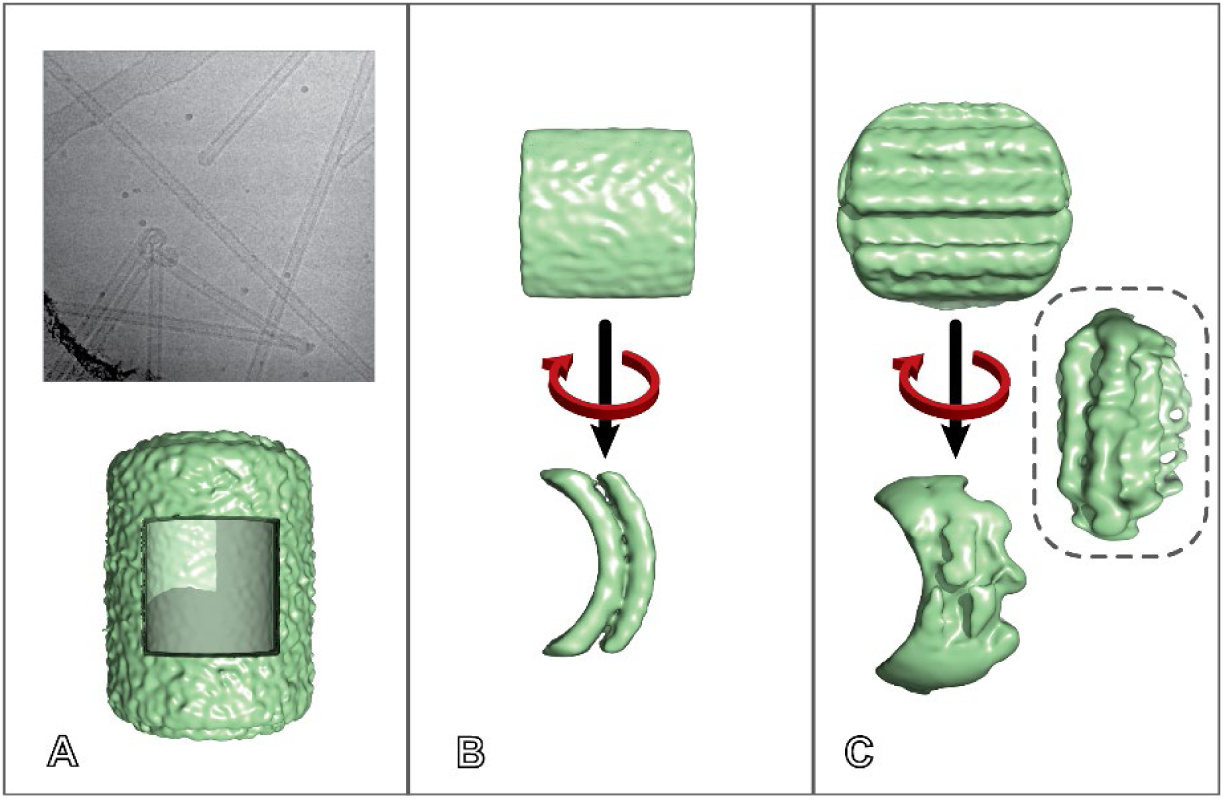
Sar1 tubules data collection and alignment. **A** Sar1 tubule micrograph (top) with azimuthal aligned average with subsection to be aligned (bottom). **B** After RASTR the subsection reconstructed using the azimuthal averaged Euler angles. **C** After alignment and classification a pattern of l inear l ines appears on the outside of the tubules, which gains definition during masking.

Despite the success in reconstructing the VipA/VipB structure using RASTR, estimating the resolution by FSC proved to be unreliable due to the masking imposed during isolation of the RASTR subparticles. FSC calculated from half-maps did not approach zero, so could not be used for reliable resolution estimation (Supp. Fig. 4A, 4B). As an alternative, we used RESMAP (Supp Fig 5,6) which reported local resolution from 3.2 to 4.2 Å. Additionally, we used map-model FSC to try and estimate the overall resolution. In the case of the VipA/VipB tubules, the map-model FSC of the upweighted sphere at 0.5 was 4.17 Å (Supp. Fig. 4C), when masked to a single VipA/VipB pair near the center of our upweighted region the FSC at 0.5 was 4.07 Å (Supp. Fig. 4D). While our reported resolution does not approach the original 3.5 Å map, we believed based on the features seen in Figures 6 and 7 that within the upweighted region the map was of similar if not slightly higher resolution than the original.

Processing of the VipA/VipB sheath using the RASTR method demonstrated that it can be used to generate 3D reconstructions of tubular particles with equivalent or better quality than helical reconstruction without imposing helical symmetry. We note, however, that we were unable to converge on the correct structure without having a good starting model. We expect this was due to inability to separate out the polarity of the helical tubule during refinement. We expect that a more sophisticated Euler search could overcome this problem. Due to the production of the azimuthal average, the Eulers for the subparticles generated by RASTR should be fairly close to their optimal angles, so searching angles of plus or minus 90° for theta and a narrow range of phi should be sufficient for a high quality reconstruction (see Supp. Fig. 1).

### Sar1…GalCer tubules

Given our success using RASTR on a helical tubule, we tried the technique on Sar1 decorated membrane tubules that have previously been shown to have local order but no long range symmetry. Previously our group showed that Sar1 oligomerizes on GalCer lipid tubules but could not be reconstructed to high resolution due to the lack of coherent long-range symmetry when it oligomerizes on membrane. We used RASTR in order to characterize the way that Sar1 oligomerizes on membrane. We collected cryo-EM data of Sar1 coated membrane tubules (Supp. Table 2). Tubes were then segmented aligned and classified with traditional 2D classification. As with our previous study, Sar1 could be observed to bind the tubules, but no long-range order was observed. We next subjected the tubules to RASTR processing. In this way, we were able to separate membrane surfaces with bare membrane from ones with Sar1 bound. Extensive 2D and 3D classification was used to separate bare tubules, unordered decorated tubules, and ordered decorated tubules. 2D classifications allowed us to see decorations and discard bare tubules. Once we knew a tubule was bare, we could discard all the particles from that specific tubule, generating a stack of decorated tubules only. Within the decorated tubules, we were able to use 3D classification to separate out ordered decorations from disordered decorations. This revealed that Sar1 oligomerized as long strings of protein that run along the flat tubular axis, with only very weak lateral interactions. The lack of lateral association was likely due to the radius of the tubules. Our previous studies showed that Sar1 made 2D lattices on relatively flat membrane but the order broke down on tubules with diameters lower than 100 nm^33^. Our new results using RASTR reveal that the reason for this was that the increased curvature of the GalCer tubes caused the lateral interactions between Sar1 to break, leaving only the vertical interactions. These observations would not have been possible without the use of RASTR.

## Conclusions

RASTR provides a new tool for processing tubules that does not require symmetry, making it more flexible than current processing methods. We anticipate that this approach will enable structure determination for a variety of biological molecules that were previously inaccessible. Two types of complexes that RASTR will be able to open new avenues for research are filament decorations and tubules with limited symmetry. Filament decorations can be examined in their natural state, bound to filaments, and their structures and binding interfaces can be explored, while limited symmetry complexes on tubules will be able to be resolved without extensive manual curation and masking. RASTR could also assist in solving the structures of filaments in which the helical symmetry is difficult to determine. While current results provide compelling evidence of the value of the technique, there are many improvements to RASTR that can be implemented. For instance, alignment and classification of RASTR particles could be improved by better subtraction that better accounts for deviation from the model and weights the model better for a cleaner subtraction. Additionally, resolution estimation requires estimating local resolution using map features instead of the typical FSC measurements. Nonetheless, RASTR has the potential to allow processing of a variety of particles (asymmetric decorations) that were previously inaccessible except through extensive manual masking and allow helical filaments to be solved without needing any knowledge of the symmetry.

## Acknowledgements

The authors acknowledge the use of instruments at the Biological Science Imaging Resource supported by Florida State University and NIH grants S10 RR025080 and S10 OD018142. The DE-64 was purchased with funds provided from NIH grant U24 GM116788 The work presented was supported by the NIH grant R01GM108753 and funds from Florida State University.

## Supplemental

### Section 1 – RASTR Arguments

- **RASTR arguments**

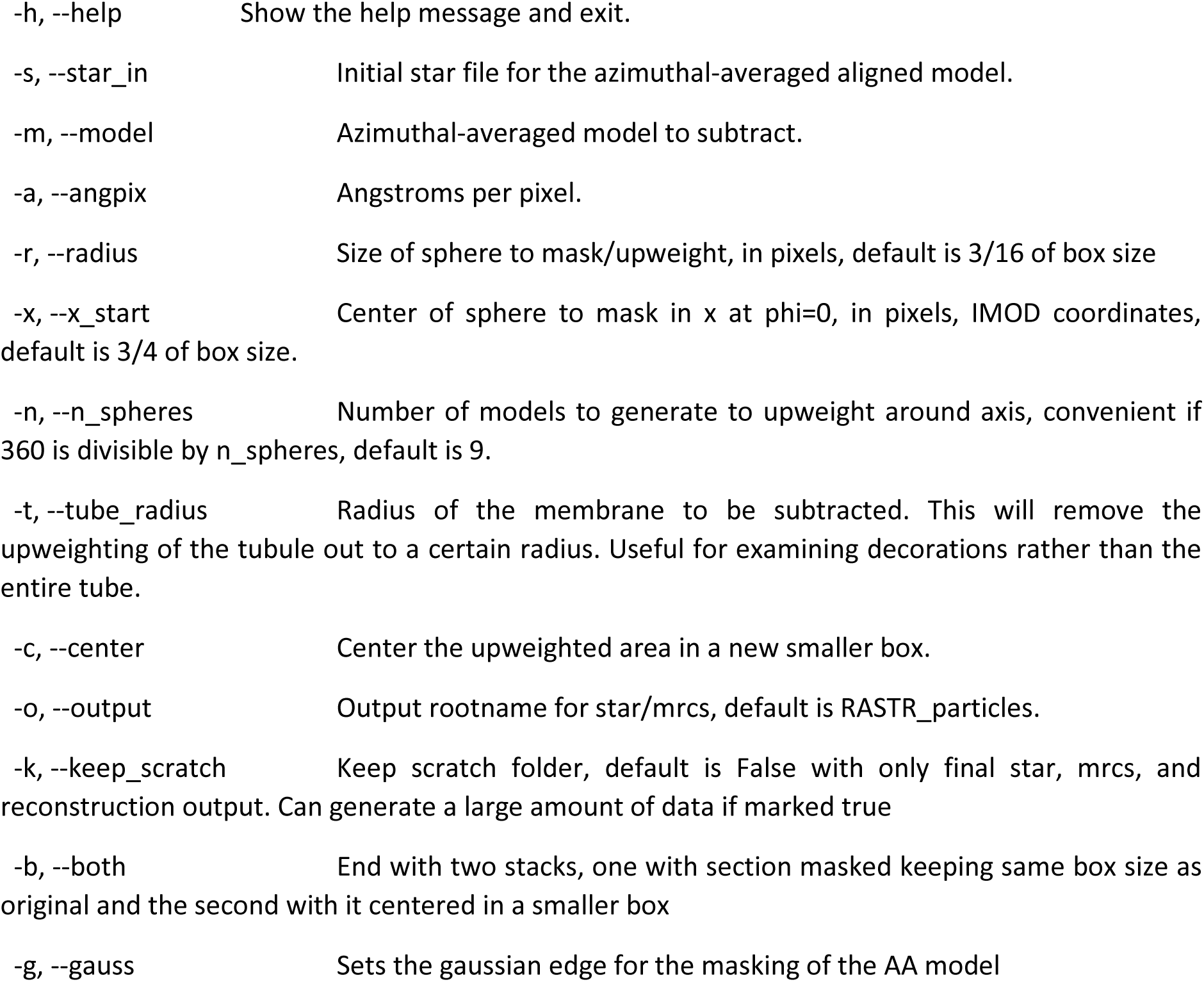

|  |  |
| --- | --- |
| -h, --help | Show the help message and exit. |
| -s, --star_in | Initial star file for the azimuthal-averaged aligned model. |
| -m, --model | Azimuthal-averaged model to subtract. |
| -a, --angpix | Angstroms per pixel. |
| -r, --radius | Size of sphere to mask/upweight, in pixels, default is 3/16 of box size |
| -x, --x_start | Center of sphere to mask in x at phi=0, in pixels, IMOD coordinates, default is 3/4 of box size. |
| -n, --n_spheres | Number of models to generate to upweight around axis, convenient if 360 is divisible by n_spheres, default is 9. |
| -t, --tube_radius | Radius of the membrane to be subtracted. This will remove the upweighting of the tubule out to a certain radius. Useful for examining decorations rather than the entire tube. |
| -c, --center | Center the upweighted area in a new smaller box. |
| -o, --output | Output rootname for star/mrcs, default is RASTR_particles. |
| -k, --keep_scratch | Keep scratch folder, default is False with only final star, mrcs, and reconstruction output. Can generate a large amount of data if marked true |
| -b, --both | End with two stacks, one with section masked keeping same box size as original and the second with it centered in a smaller box |
| -g, --gauss | Sets the gaussian edge for the masking of the AA model |

### Section 2 – Data Collection Statistics

**Supp. Table 1.**
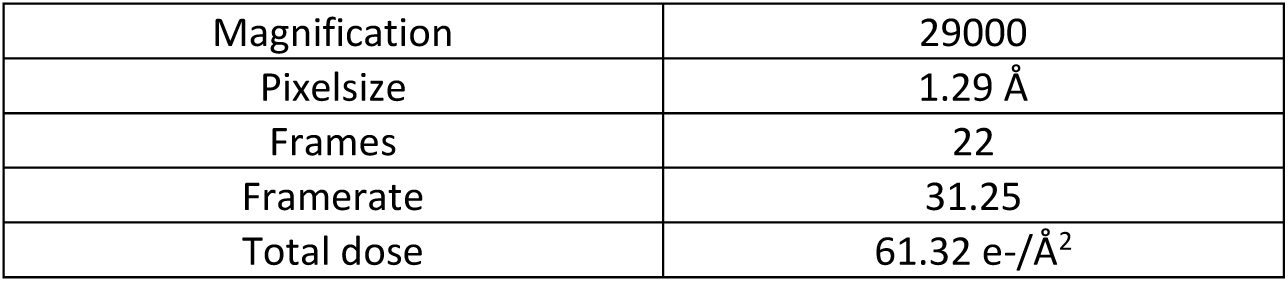
Collection Statistics for undecorated GalCer tubes.

**Supp. Table 2.**
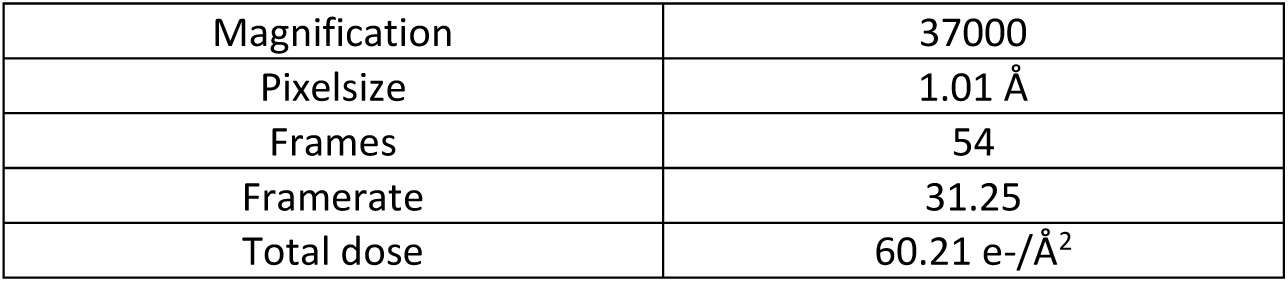
Collection Statistics for Sar 1…Gal Cer tubes.

### Section 3 – Euler Distribution

**S Figure 1.**
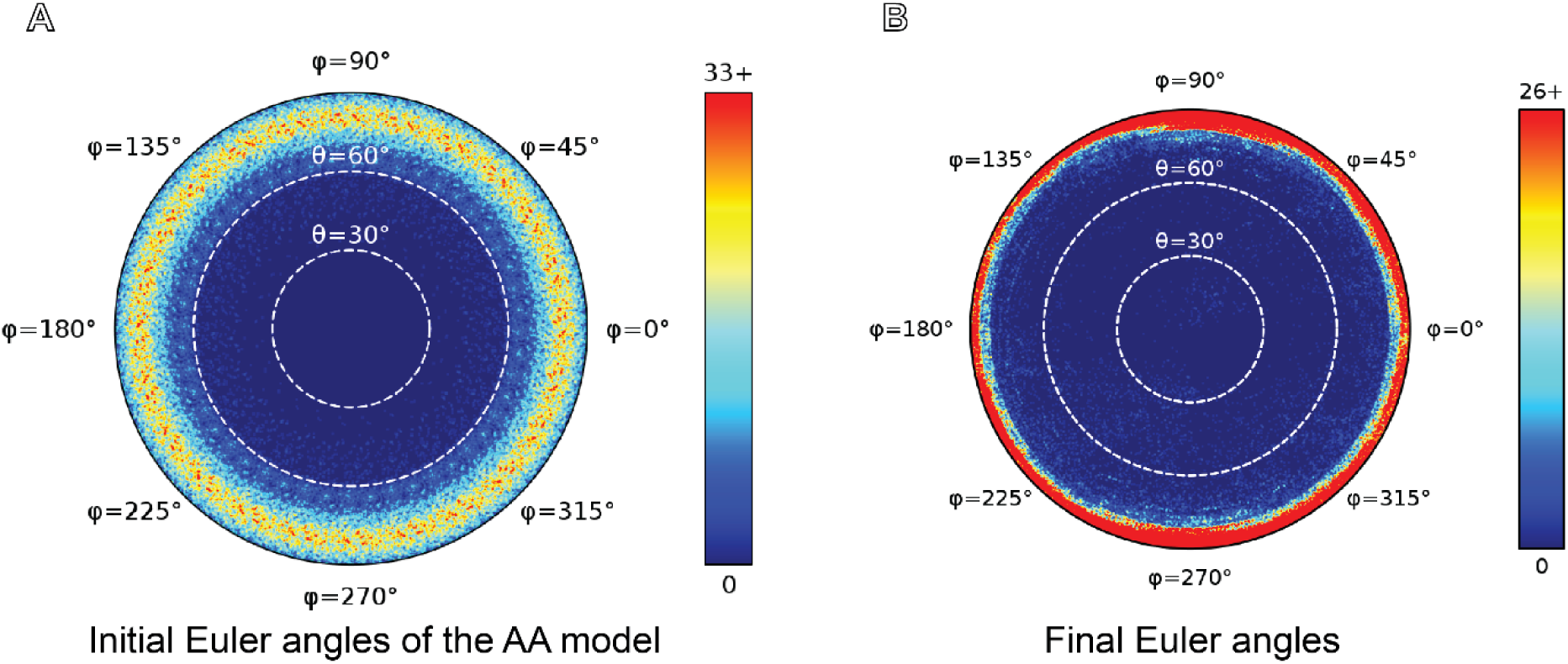
VipA/VipB Euler Angles. A Initial randomized Eulers for the AA model shows an even distribution. **B** Final Euler angles demonstrate a similar distribution of phi.

### Section 4 – ResMap and Model-Map FSC VipA/VipB

**S. Figure 2.**
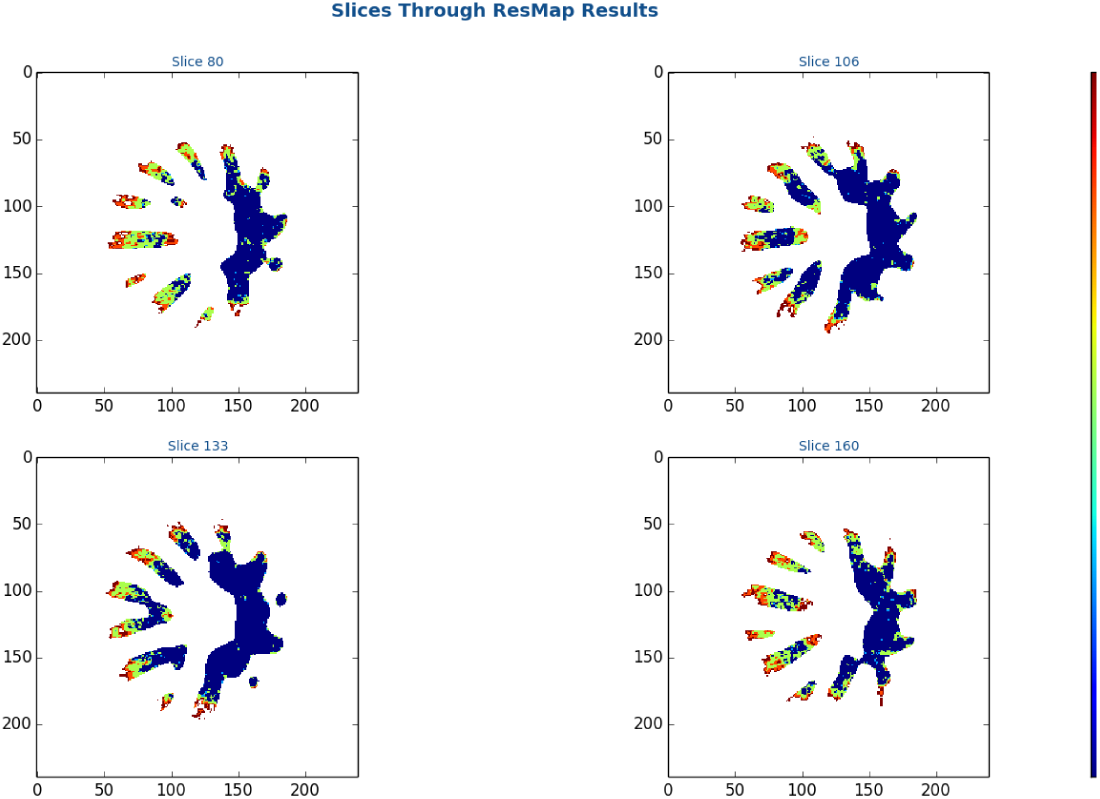
ResMap local resolution slices for ideal Vip A/Vip B. Within our final map (pre-sharpening) local resolution was reported between 4. 4 and 8. 4 Å.

**S. Figure 3.**
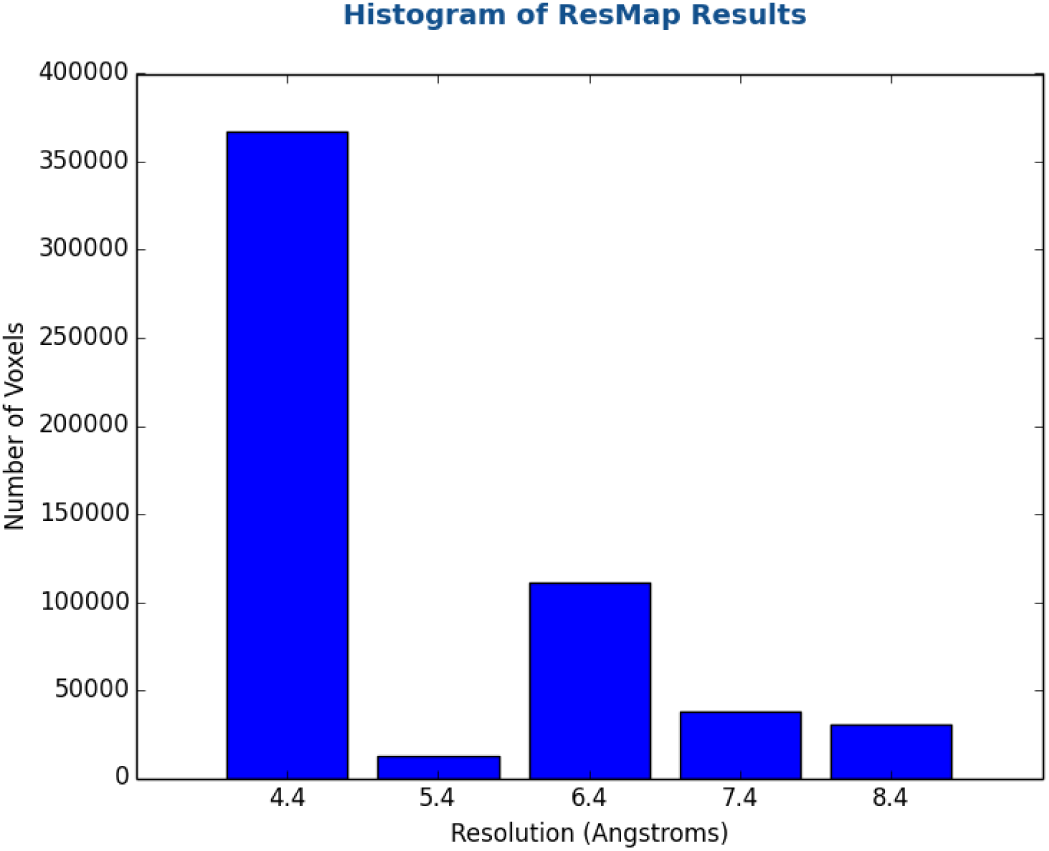
ResMap local resolution histogram of ideal Vip A/Vip B. Within our final map (pre-sharpening) local resolution was reported between 4. 4 and 8. 4 Å.

**S. Figure 4.**
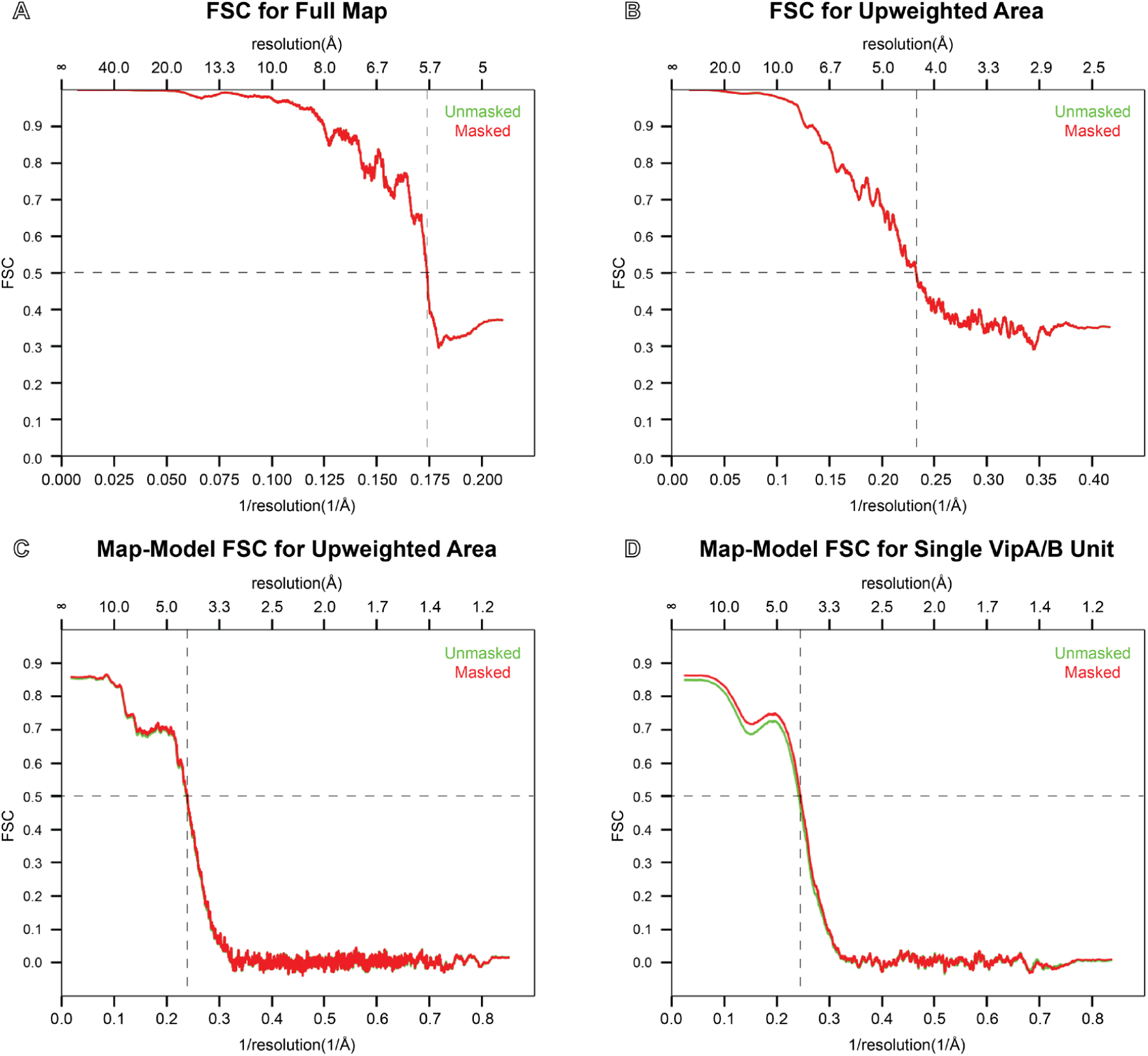
FSC calculations for RASTR processed experimental Vip A/Vip B experimental structure. Calculated in Phenix Mtriage. **A** Half-map FSC on the VipA/ VipB full map including downweighted area (0. 5 Res = 5. 7 Å). **B** Half-map FSC of the VipA/ VipB map clipped to RASTR upweighted section (0. 5 Res = 4. 3 Å). **C** Map-Model FSC on the VipA/ VipB map clipped to upweighted section (0. 5 Res = 4. 17 Å). **D** Map-Model FSC on a single VipA/ VipB subunit (0. 5 Res = 4. 07 Å).

**S. Figure 5.**
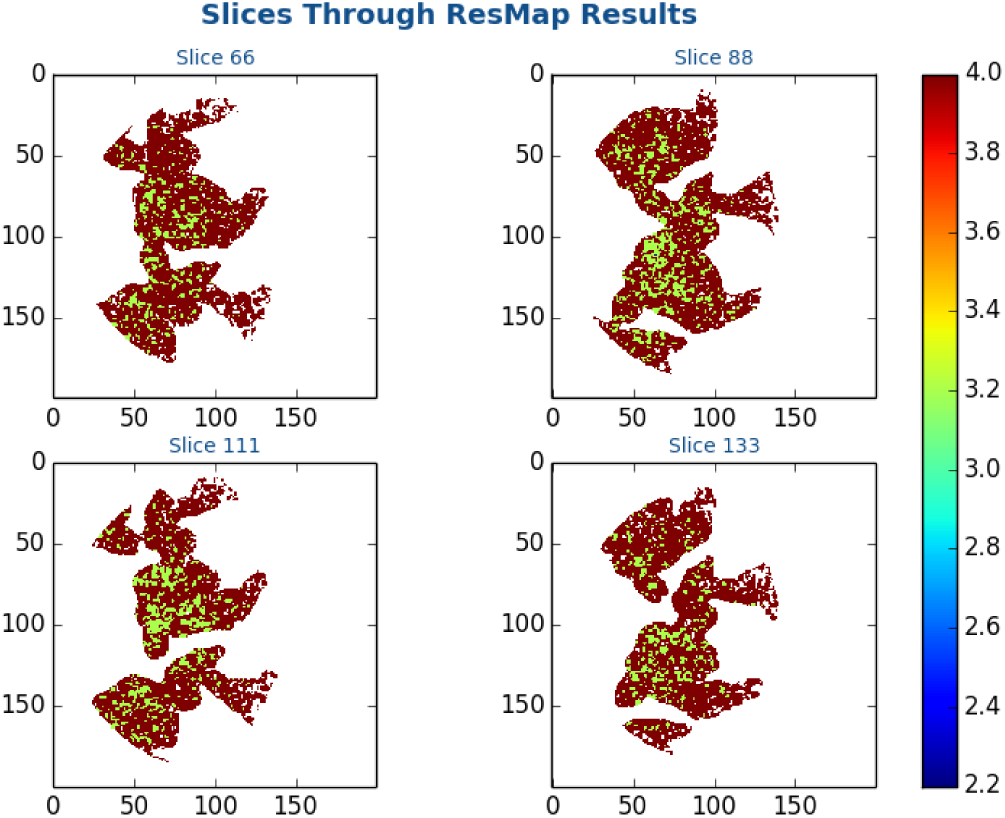
ResMap local resolution slices of experimental Vip A/Vip B. Within our final map (pre-sharpening) local resolution was reported between 3. 2 and 4.2 Å.

**S. Figure 6.**
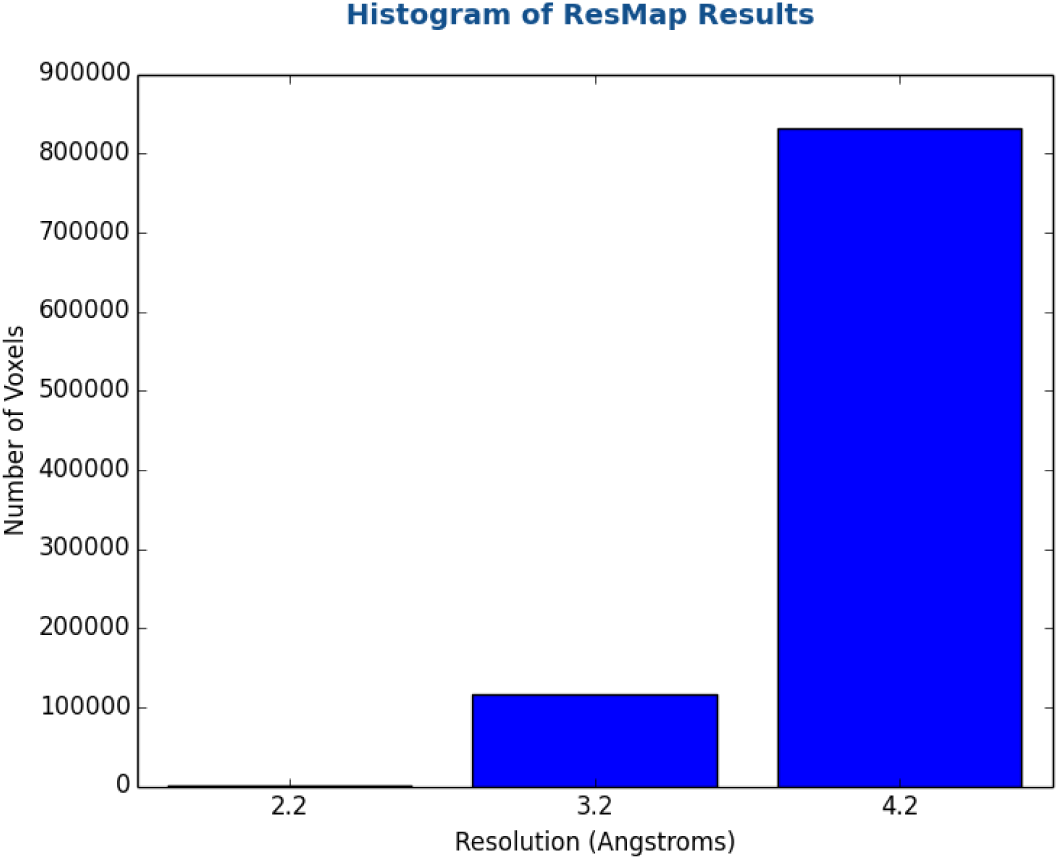
ResMap local resolution histogram of experimental Vip A/Vip B. Within our final map (pre-sharpening) local resolution was reported between 3. 2 and 4. 2 Å.

